# Exploring Differential Gene Expression Profiles Of *Dero (Allodero) Hylae* In Their Parasitic And Free-Living Forms

**DOI:** 10.1101/2022.07.17.500331

**Authors:** Claire Bonham, Ashley Roguski, Gabriel Langford, Jason Macrander

**Author notes:** These authors contributed equally to this work (Claire Bonham, Ashley Roguski).

## Abstract

Parasitism is ubiquitous in nature, yet little is known about the evolutionary mechanisms that lead to a parasitic lifestyle. Facultative parasites can switch between free-living and parasitic lifestyles, which may provide an opportunity to study the genetic mechanisms underlying a transition to parasitism. The oligochaete *Dero (Allodero) hylae* is a facultative parasite commonly found within the ureter of various anuran species, such as the Cuban Tree Frog (*Osteopilus septentrionalis*). *Dero hylae* makes passage through the frog’s cloaca, where it then infects the ureter. In the ureter, the worm loses free-living characteristics such as hair setae, dorsal setae, a digestive tract, and fossa with gills as it transitions to a parasitic lifestyle. *Dero hylae* may be expelled from its host during urination, when this occurs the worm will reacquire free-living characteristics. The focus of this study is to compare the differential gene expression profiles observed when this rapid morphological change takes place. Specimens of *D. hylae* were collected from wild Cuban Tree Frogs and either flash-frozen for their parasitic stage RNA profile or cultured for two weeks to produce their free-living stage and then flash-frozen. Using the sequenced RNA, a *de novo* transcriptome was assembled and differential gene expression RNA Tag-Seq analysis between the free-living and parasitic life forms was analyzed. Based on these results, we have identified 213 genes differentially expressed transcripts between the two life forms, 190 of these being up-regulated in the free-living life form. While over half of the differential genes recovered did not recover any significant BLAST hits, many of these genes did provide insight into which molecular signals are potentially used by *D. hylae* to lose and subsequently regrow their setae, digestive tract, and gills. This analysis provides significant insight into which differentially expressed genes are linked to drastic morphological changes observed in this rare oligochaete parasitism across the free-living and parasitic forms of *D. hylae*.

## INTRODUCTION

Parasitism has evolved independently on numerous occasions (Roberts et al., 2012), however, we understand little about the adaptive process that leads to a parasitic lifestyle (Dieterich and Sommer, 2009; Viney, 2017; Luong and Mathot, 2019). As reviewed by Luong and Mathot (2019), facultative parasites may provide a roadmap to understanding obligate parasitism. Facultative parasites can transition between free-living and parasitic lifestyles under specific environmental conditions, providing an opportunity to understand the genetic mechanisms that facilitate a transition to a parasitic lifestyle. The phenotypically plastic annelids in the genus *Dero* and their anuran hosts may serve as a model system to study the evolutionary transition to parasitism (Gelder, 1980; Andrews et al., 2015). While few studies have been conducted on these rare endoparasitic annelids, it is well-established that *Dero* worms facultatively parasitize or phoretically travel on anuran hosts (Lopez et al., 1999; Andrews et al., 2015). It is unknown what genetic mechanisms facilitated the presumed progression from phoresy to facultative parasitism in some species of *Dero*.

Michaelson (1926) identified the first oligochaete symbiont in the ureter of South American hylid tree frogs. These collected specimens lacked dorsal setae, branchial fossa, and gills, which are characteristics of almost all oligochaetes. However, once the worms were cultured in pond water, they developed dorsal setae, caudal fossa, and gills (Michaelson, 1926). Based on the unique parasitic lifestyle and morphological characteristics of these worms, the subgenus *Allodero* was created and placed in the genus *Dero* based on the formation of the caudal fossa and gills when free-living (Sperber, 1948). Within *Dero (Allodero)*, there are 7 known species that infect either the eyes or the ureters of anurans (Sinch et al., 2019). Sinch et al. (2019) used molecular evidence to conclude that the subgenera in *Dero: Allodero, Aulophorus*, and *Dero* are not distinct evolutionary lineages and that the genus is paraphyletic because it includes *Branchiodrilus*.

The parasitic oligochaete *Dero (Allodero) hylae* is found within anuran species across tropical and subtropical regions (Harman, 1973; Harman & Lawler, 1975). *Dero hylae* is only found within the families Hylidae and Bufonidae (Harman and Lawler, 1975). The host of *D. hylae* seems to be dependent on the region, as, in the Southeastern Coastal Plain of the United States, the worms have been noted to infect *Hyla cinerea, Hyla versicolor*, and *Hyla squirella* (Harman and Lawler, 1975). Meanwhile, in Louisiana, *H. cinerea* is not known to be infected while *H. versicolor* and *H. squirella* are known hosts (Harman, 1973).

*Dero hylae* is capable of locating and infecting new hosts via the hosts’ urine, as it travels into the ureter by passage of the cloaca, where the worm adopts an endoparasitic lifestyle (Harman and Lawler, 1975; Andrews et al., 2015). Within 72 hours of a free-living worm infecting a host, they begin to lose their hair setae, some of their dorsal setae, and gills in the fossa are minimized (Fig.1). It was previously thought that *D. hylae* did not cause the ureter any harm or damage (Harman and Lawler, 1975), which sparked debate on the nature of the relationship between anurans and *D. hylae*. However, it has recently been experimentally demonstrated that infections cause physical damage to the ureter that can kill young frogs (Andrews et al., 2015).

**Figure 1:**
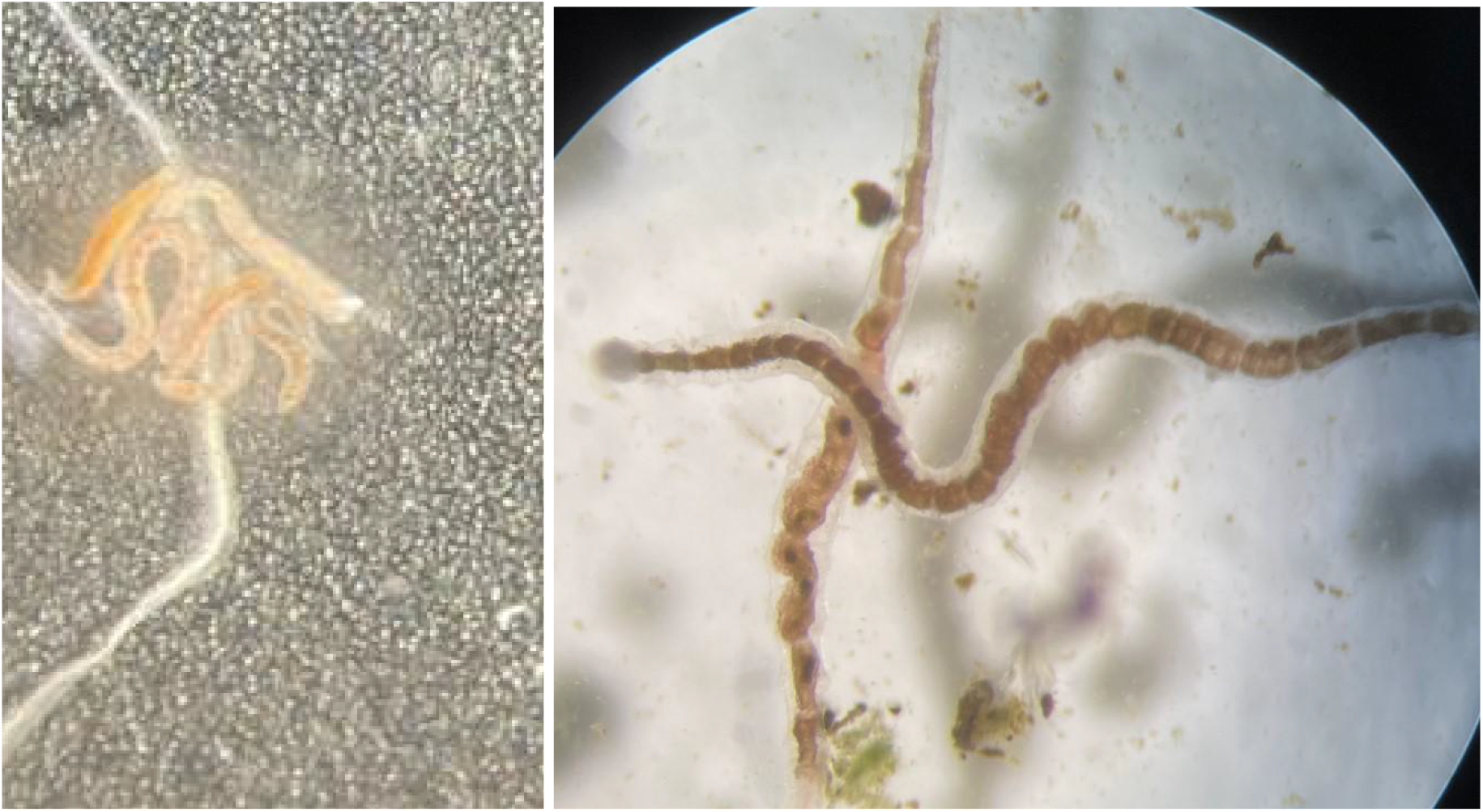
Photos depicting the morphological transformation D. hylae go through from their parasitic (left) to free-living (right) distinct morphological life-history stages.

Andrews et al. (2015) also demonstrated that parasitic worms could be expelled from their host during urination. During intense infections, worms effectively spill over from the ureter into the urinary bladder and are susceptible to being voided with the urine. If they do not find a new host after 72 hours or more after expulsion, *D. hylae* will begin a morphological transformation to grow hair setae, dorsal setae, a digestive tract, and a fossa with gills. This major morphological transformation provides the previously parasitic worms with the necessary anatomical structures of their free-living environment. The worms are capable of surviving in a free-living environment for an extended time period, although it is unclear under what circumstances free-living *D. hylae* seek anuran hosts (Andrews et al., 2015).

It is currently unknown how these worms reproduce, and whether or not the free-living form is required for reproduction (Lutz, 1927; Andrews et al., 2015). However, some studies have suggested that worms may only reproduce asexually while in the host, which would require the worms to leave the host to infect another (Goodchild, 1951; Gelder, 1980). Andrews et al. (2015) suggested that once acclimated to a free-living environment *D. hylae* may develop sexual organs under specific environmental conditions, however, they were unable to stimulate development of sexual organs under laboratory conditions.

The mechanism of transition between free-living to parasitic and vice versa is also unknown, with a lack of genetic material among these parasitic oligochaetes to aid our understanding. By collecting *D. hylae* in their parasitic and free-living forms RNA could be extracted and differential gene expression quantified at each life-history stage. We aim to determine which genes may better assist our understanding of the morphological transformation between the free-living or parasitic forms in *D. hylae*.

## METHODS

### Sample collection

This study was performed on the campus of Florida Southern College in Lakeland, Florida, USA. From which, wild Cuban tree frogs (*Osteopilus septentrionalis*) were caught by hand, or net if necessary, to obtain samples of *D. hylae*. The frogs were transported to the laboratory and promptly euthanized by double pithing, in accordance with IACUC protocol #2021-03. Following euthanization, the urinary bladder and ureter of the Cuban tree frogs were dissected to collect parasitic *D. hylae* samples. All *D. hylae* samples found within the frogs were either flash-frozen with liquid nitrogen for later RNA extraction or placed in a small dish of water from the campus mesocosms to culture free-living forms. Worms were observed daily for the development of their setae, gut, and gills to determine when the morphological transformation had occurred from their parasitic to free-living forms. Once in their free-living form, they were flash-frozen in liquid nitrogen for RNA extraction. Each sample had approximately 6 worms of each corresponding life history stage. These were collected in replicates of 4 each, with the three best RNA extractions sent out for sequencing.

### RNA extraction and sequencing

RNA extractions were completed using the Invitrogen RNAqueous™ extraction kit (Thermofisher Scientific, Waltham, MA, USA) following the manufacturer’s protocol. RNA concentrations were measured on the NanoDrop ND-1000 Spectrophotometer (ThermoFisher) with the three highest concentrations for each morphological state sent to Admera Health Biopharma Services (South Plainfield, NJ, United States) to be sequenced. RNA aliquots from all high concentration samples were combined for total RNA sequencing. At Admera Health, each of the individual and combined RNA samples’ quality and quantity were evaluated using the Qubit RNA HS assay (ThermoFisher) and Bioanalyzer 2100 Eukaryote Total RNA Nano (Agilent Technologies, CA, USA), respectively. After passing through quality control, the combined RNA samples were sequenced using the NEBNext^®^ Ultra™ II Non-Directional RNA Second Strand Synthesis Module (New England BioLab, MA, USA) on the Illumina NovaSeq - S4 flow cell (CA, USA), at a minimum of 40 million paired end reads across 150 base pairs in both directions. The individual RNA Tag-Seq samples were prepared for RNA Tag-Seq using the QuantSeq 3’ mRNA-Seq Library Prep Kit FWD for Illumina (Lexogen, Vienna, Austria) at a minimum of 100 bases and 4 million reads per sample. Raw reads were deposited on NCBI’s Sequence Read Archive (SRA) database under the BioProject accession PRJNA854921.

### Transcriptome assembly and gene expression analysis

We conducted a de novo transcriptome assembly for *D. hylae* using Trinity v2.13.2 (Grabherr et al. 2011) using Trimmomatic (Bolger et al. 2014) and the “no_bowtie” option, with all other default parameters (Haas et al. 2013). Transcriptome completeness was determined using the metazoan database in the program BUSCO v5.3.2 (Manni et al. 2021). Once assembled, raw reads were mapped to the assembled transcriptome using Kallisto (Bray, et al., 2016) to generate transcript and gene abundance measures. A differential expression analysis comparing RNA Taq-Seq runs across the two distinct morphological life history stages was conducted using EdgeR with a fold change of 4 or greater and 0.05 P-value for significance (Robinson, et al., 2010). Differentially expressed genes were further screened through subsequent tBLASTn searches (Altschul, et al., 1990) against the annotated UniProt dataset (Uniprot Consortium, 2018) and non-redundant NCBI protein database to predict gene function and identify gene ontology groups to aid in our understanding of these gene functions as they relate to the morphological transformation between parasitic and free-living forms.

## RESULTS

In total, there were 17 Cuban Tree Frogs collected from FSC campus and surrounding areas from November 2021 to January 2022. Of these, only two were found to be infected with *D. hylae* (11.7%). Although lower than expected, it does coincide with a lower prevalence previously surveyed with colder months (Andrews, et al., 2015). Each contained a large number of parasitic *D. hylae* worms, however, it took approximately two weeks for culturing of *D. hylae* to induce the morphological transformation to their free-living form to display setae and a digestive tract, which differs from previous reports of approximately 72 hours (Andrews, et al., 2015), which again may be attributed to slower life cycle stages during the colder months. Despite sampling only two frogs, there were several *D. hylae* from each to collect and use for subsequent RNA extractions.

For the transcriptome assembly 21.7 million paired end reads were sequenced and screened for quality control, of which 17.2 million were used to assemble 91,486 trinity ‘genes’ and 100,319 transcripts, with a contig N50 of 787 bases. The BUSCO completeness was 74.5% based on the metazoan database, which was on par with previously assembled parasitic transcriptomes using similar approaches (Langeland et al. 2021; Ahmed et al 2021). The number of raw reads across Tag-Seq samples ranged from 2.4 to 2.7 million raw reads, with 42 - 56% of each sample pseudo-aligning within kallisto to the assembled transcriptome.

Our differential gene expression analysis identified 213 differentially expressed genes between the free-living and parasitic life forms. 190 of these genes were up-regulated in the free-living life form, while 23 were down-regulated. After analyzing the results of the BLAST searches of these genes we linked hits to predicted gene function and determined that over half of the differentially expressed genes did not recover any significant hits (Fig. 2). Of those with possible gene functions, multiple were solely up-regulated in the free-living life form and further characterized based on Gene Ontology annotations associated with the molecular function and biological processes (Table 1), with multiple genes involved in actin binding, metal ion binding, ATP binding, or calcium ion binding. Aside from the Gene Ontology analysis, we found several genes of interest based on protein descriptions that were upregulated in the free-living life form and one in the parasitic life form. These genes have varying functions related to the morphological transition from parasite to free-living (Table 2).

**Figure 2:**
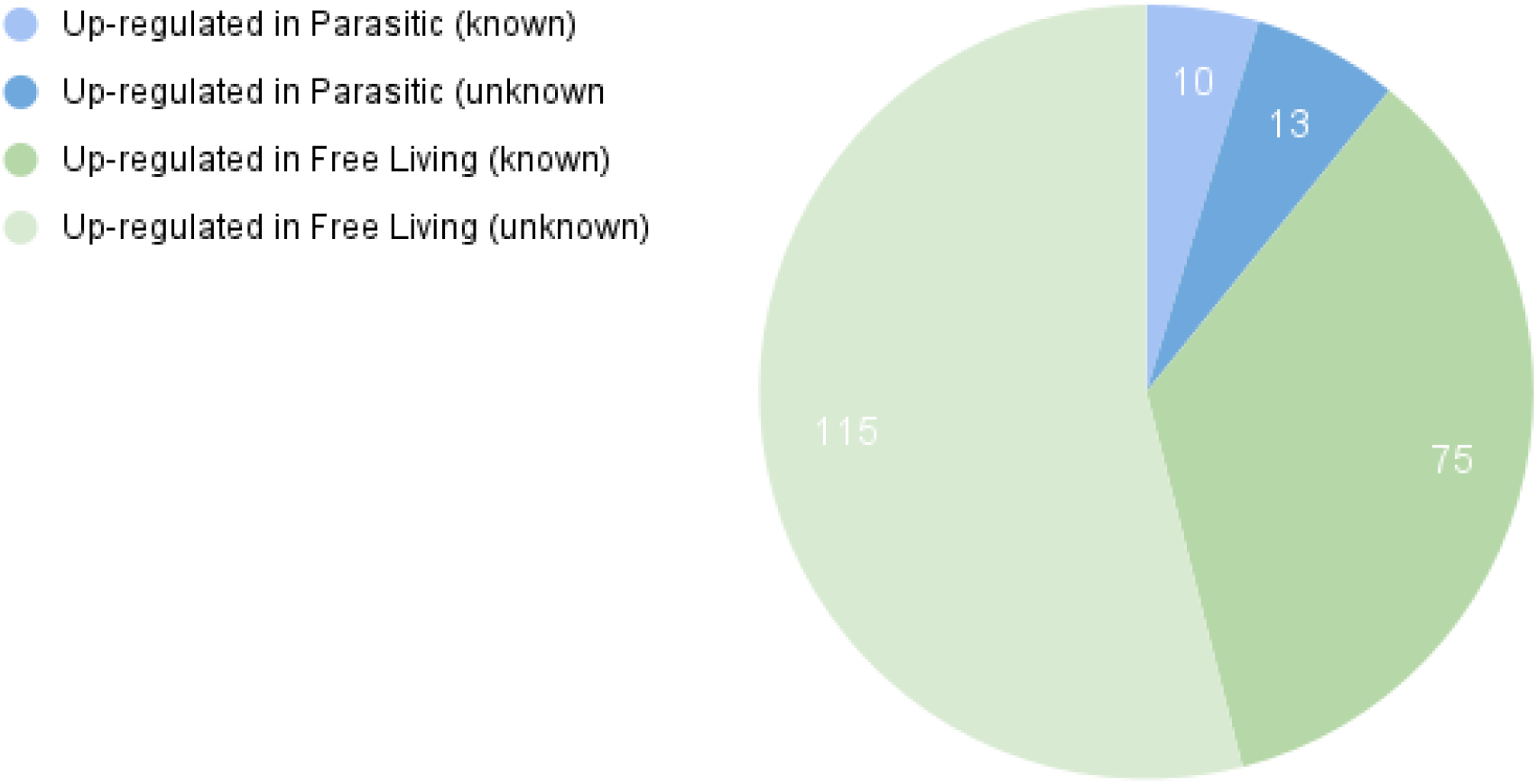
Pie chart displaying the identified genes previously annotated by past research in both the free-living and parasitic groups. Parasitic genes are denoted by light blue (known descriptions of gene function) and dark blue (unknown gene function) while free-living genes are denoted by dark green (known description of gene function) and light green (unknown description of gene function).

**Table 1:**
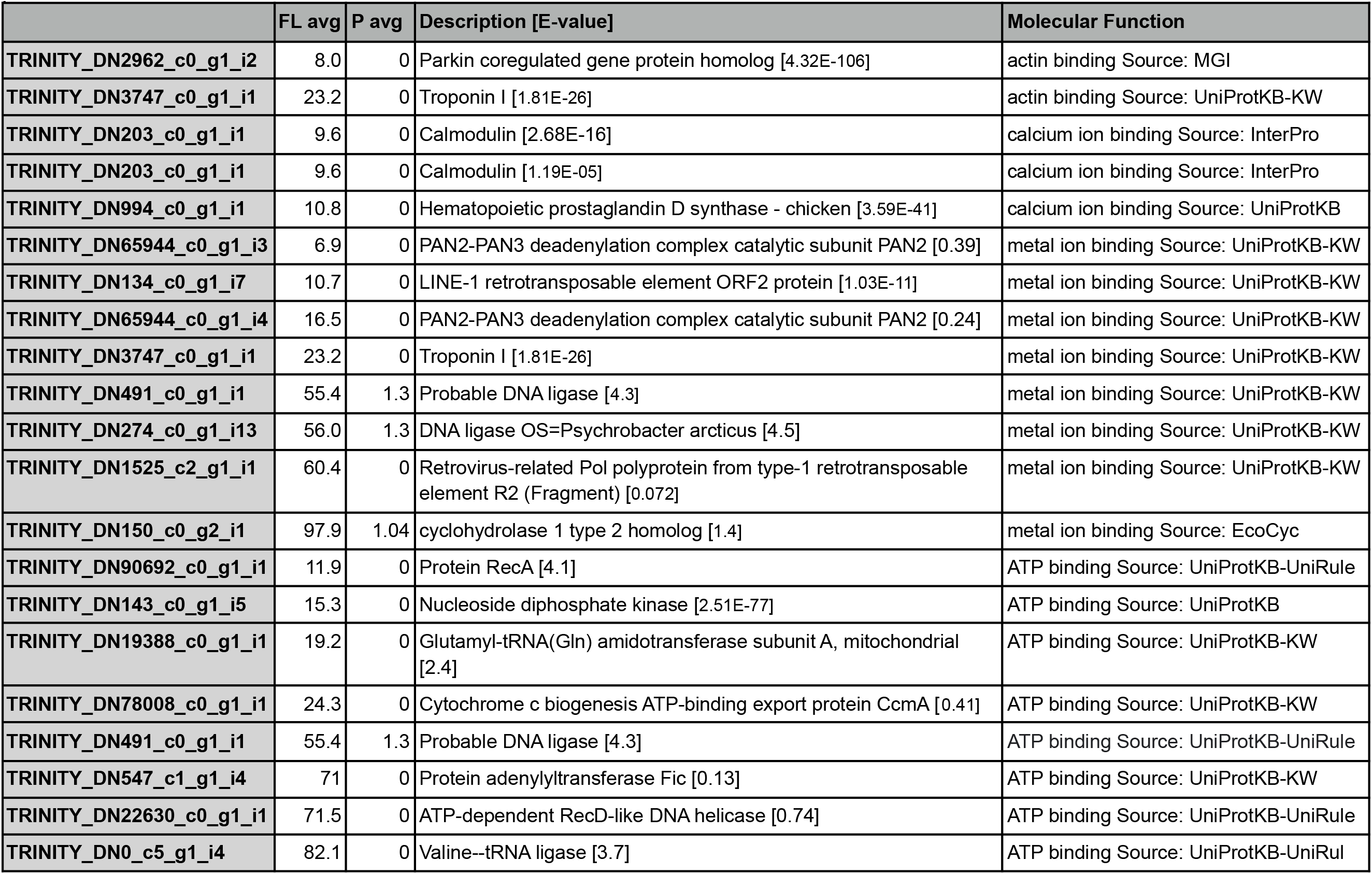
Actin, Calcium, Metal Ion, and ATP Binding related genes and their average expression value (TPM) in free-living and parasitic life forms.

**Table 2:** Genes of notable function and their average TPM in both free-living and parasitic life forms.

|  | FL avg | P avg | Description | FUNCTION - GO MOLECULAR |
| --- | --- | --- | --- | --- |
| TRINITY_DN66664_c0_g1_i1 | 152.8 | 1.2 | Bombyxin B-2 | growth factor activity Source: InterPro<br>hormone activity Source: UniProtKB-KW |
| TRINITY_DN492_c0_g1_i2 | 55.8 | 0 | Caveolin-1 | caveola assembly Source: InterPro<br>T cell costimulation Source: UniProtKB |
| TRINITY_DN10_c2_g1_i1 | 16571.8 | 38.1 | Protein TAR1 | Regulation of cellular respiration |

## DISCUSSION

To our knowledge this is the first comparative transcriptomic analysis from an endoparasitic annelid that exhibits such a sharp contrast in morphology in both their free-living and parasitic life stages. Our analysis indicates asexual reproduction is highly likely as only 213 differentially expressed genes were recovered from over one hundred thousand transcripts, however, it is still unknown whether or not *D. hylae* ever reproduces sexually. These results indicate that gene functions associated with these candidate transcripts likely play a key role during this morphological transformation. Additionally, there was a large number of genes that lacked any significant BLAST matches, indicating that there is much to learn about the transition this parasite takes on multiple times throughout its life cycle as there are limited genomic resources available to study this process. The functions for many of the genes that were described had a wide variety of Gene Ontology designations within the Biological Processes and Molecular Function domain, however, some patterns emerged that may link these genes to the switch from free-living to parasitic.

Of the up-regulated genes in the free-living form, we found that several were linked to rapid morphological change or development. One gene in particular, BBXB2 (TRINITY_DN66664_c0_g1_i1) produces the protein Bombyxin B-2, which regulates metabolism and stimulates tissue growth in insects (Kawabe et al., 2019). Bombyxin has also been shown to activate the production of ecdysone in insects by stimulating the prothoracic gland. The production of ecdysone causes larval insects to begin their development into their adult stage (Iwami, 2000). While *D. hylae* is no insect, the presence of this gene that has been linked to developmental changes and movements through the life cycle in other organisms indicates that it likely has a shared function here and is potentially involved with signaling to *D. hylae* when to make the switch from parasite to free-living.

Another gene that was up-regulated in the free-living life form of *D. hylae* was Cav-1, which forms the protein Caveolin-1 (Table 2) This protein has been researched heavily as it is associated with many different forms of cancer. Caveolin-1 regulates cellular metabolism, and in cancer can influence tumor growth by halting apoptosis and encouraging metastasis (Nwosu et al., 2016). Previous studies have confirmed Cav-1s involvement in cellular proliferation, which makes sense in the context of our study as Cav-1 was up-regulated in free-living *D. hylae*, suggesting that the presence of Cav-1 contributed to the growth of setae, digestive tract, and caudal fossa within our samples (Boscher and Nabi, 2012).

There were 7 genes up-regulated in the free-living life form that was associated with metal ion binding (Table 1). Specifically: Pan2, LINE-1 retrotransposable element ORF2 protein, Troponin I, lig, ligA, Retrovirus-related Pol polyprotein from type-1 retrotransposable element R2, and TC_0384 which is involved in GTP cyclohydrolase 1 type 2 homolog. None of the genes up-regulated in the parasitic life form were associated with metal ion binding. This is relevant as metal ion binding plays an important role in directing different processes such as cellular differentiation (Leszczynski and Shukla, 2014). The presence of genes involved in metal ion binding only within the free-living life form leads us to hypothesize that their involvement in cellular differentiation aids *D. hylae* in the regrowth of free-living traits.

Further, we found multiple genes involved in ATP binding up-regulated in the free-living parasite samples (Table 1). ATP plays a crucial role in a number of biological pathways and mechanisms such as signal regulations, DNA-binding, and enzyme activity. Specifically, ATP-binding is involved in a number of metabolic processes as well as cellular motility, muscle contraction, and membrane transport (Chauhan, 2009). Recent studies have found that ATP-binding is involved in the translation or regulation of gene expression, DNA repair, and serves as a physiological function in multiple protozoan parasites (Sauvage et al., 2009). In *D. hylae*, eight genes associated with ATP-binding were up-regulated in the free-living form and one gene associated with ATP-binding was up-regulated in the parasitic form. Although ATP and ATP-binding have a variety of functions and involvement in cellular processes, their abundance found in the free-living form of *D. hylae* is significant. The lack of ATP binding genes associated with the parasitic form of *D. hylae* could suggest that the parasitic form does not require as many mechanisms and pathways as the free-living form, resulting in a decreased amount of cell differentiation and protein function. This large variation in genetic expression between free-living and parasitic forms of *D. hylae* guides us to the idea that the presence of these genes is associated with cellular differentiation when the regrowth of free-living traits occurs.

Four genes that were up-regulated in the free-living *D. hylae* were associated with calcium ion binding (Table 1). Calcium ions and calcium bonding proteins are necessary to maintain homeostasis and regulate numerous cellular functions. Previous studies have determined that calcium ions are essential for host cell invasion in parasites that have an intracellular life cycle, such as trypanosomatids (Docampo and Huang, 2015). It has also been found that calcium ions function to regulate cell bioenergetics and aid in sensing the environment in trypanosomatids (Docampo and Huang, 2015). Calcium-dependent protein kinases and calcium-mediated signaling are involved in a number of vital functions in apicomplexan parasites, such as cell invasion, cell differentiation, protein secretion, and motility (Nagamune et al., 2008). The association with calcium ion binding found in the free-living *D. hylae*, but not the parasitic *D. hylae*, suggests that the free-living form of the parasite has a larger variety in cell differentiation and function which could play a role in the morphological changes that occur.

We also found genes associated with actin-binding were highly up-regulated in the free-living form, while no genes in the parasitic form were associated with actin-binding (Table 1). Previous literature has described how actin plays a role in other parasitic species. Some parasites, such as *Toxoplasma gondii*, rely on the actin cytoskeleton to enter host cells while some apicomplexan parasites, such as *Plasmodium*, employ their own actomyosin motor proteins to enter host cells (Axisa et al., 2000). Understanding actin-binding and actin modulating proteins in parasites are of great interest because it may suggest that actin-binding genes in the free-living form of *D. hylae* are associated with cell movement and intracellular transportation, which could be assisting in the growth of free-living traits.

Even further, three genes that encode for protein TAR1 were up-regulated in the free-living form and one was up-regulated in the parasitic form of *D.hylae*. Multiple of the identified genes encoded for protein TAR1 may be involved in the stabilization of mtDNA, the regulation of mitochondrial gene expression, and the regulation of cellular respiration. Overexpression of protein TAR1 has been determined to inhibit a respiration-deficient phenotype in mitochondrial RNA polymerase in *Saccharomyces cerevisiae* (Coelho et al. 2002). This suppression impacts the stability of mtDNA and mitochondrial gene expression. In addition, other studies have identified a significant difference in the abundance and expression of protein TAR1 in other parasitic species’ life cycles. In the tick parasite, *Theileria annulata*, it was found that protein TAR1 is increased in the transcript levels during the differentiation of the parasite to the merozoite stage but decreased in the piroplasm stage (Ingram, 1997). While the life cycle of *D.hylae* remains unknown, it can be suggested that protein TAR1 may play a role in the transformation of *D. hylae* due to the increased expression level found in the free-living form.

Two genes that were up-regulated in the parasitic *D. hylae* were associated with ribosomal proteins S6 and S8. In a study on *Plasmodium* parasites, ribosomal protein S6 was determined to regulate cell growth and the availability of glucose and lipids, which may aid in the regulation of *Plasmodium* infection. It was suggested that *Plasmodium* parasites may be selecting a favorable environment through selection or manipulation when developing within a cell with high phosphorylation of ribosomal protein S6 (Glennon et al., 2019). The up-regulated expression of ribosomal protein S6 may benefit the species in the parasitic form similarly to ribosomal protein S6 in *Plasmodium* parasites.

While it is unknown exactly how all of these genes we identified within the two life forms are related to the morphological transformation *D. hylae* undertakes, the discovery of the extreme difference in genetic expression is a good first step to learning more about this process and facultative parasites that go through similar transformations. We have uncovered multiple genes that may be linked to this morphological change, however, over half of the genes we found were not previously described, so it is likely these unknown genes also play a major role in this transformation. In the future, as these genes are described we may be able to discern more about how these genes correlate to the drastic morphological transformation in *D. hylae* and other parasites.

## ACKNOWLEDGMENTS

This research was conducted at Florida Southern College under the Institutional Animal Care and Use Committee (IACUC) protocol # 2021-03. We thank FSC’s Biology Department for funding this study and students enrolled in FSC’s capstone research class for their valuable feedback on this project and manuscript.

## References

Ahmed, M., Adedidran, F., and O. Holovachov. 2021. A Draft Transcriptome Of A Parasite *Neocamacolaimus parasiticus* (Camacolaimidae, Plectida). Journal of Nematology 53: 1–4.

Altschul, S.F., Gish, W., Miller, W., Myers, E.W. and D.J. Lipman. 1990. Basic Local Alignment Search Tool. Journal of Molecular Biology 215:403–410.

Andrews, J. M., Childress, J. N., Iakovidis, T. J., and G.J. Langford. 2015. Elucidating The Life History And Ecological Aspects Of Allodero Hylae(Annelida: Clitellata: Naididae), A Parasitic Oligochaete Of Invasive Cuban Tree Frogs In Florida. Journal Of Parasitology, 101, 275–281.

Axisa, S., Boleti, H., Couture-Tosi, E., Poupeland, C., And I. Tardieux. 2000. Molecular Biology Of The Cell. Toxofilin, A Novel Actin-Binding Protein From Toxoplasma Gondii, Sequesters Actin Monomers And Caps Actin Filaments. 11:1, 355–368.

Bolger, A. M., Lohse, M., And B. Usadel. 2014. Trimmomatic: A Flexible Trimmer For Illumina Sequence Data. Bioinformatics, 30, 2114–2120.

Boscher, C., And I.R. Nabi. 2012. Caveolin-1: Role In Cell Signaling. In: Jasmin, Jf., Frank, P.G., Lisanti, M.P. (Eds) Caveolins And Caveolae. Advances In Experimental Medicine And Biology, Vol 729. Springer, New York, Ny.

Bray, N.L., Pimentel, H., And Pachter, P. M. And L. Patcher. 2016. Near-Optimal Probabilistic Rna-Seq Quantification. Nature Biotechnology. 34(5): 525–527.

Chauhan, J.S., Mishra, N.K., And G.P. Raghava. 2009. Identification Of Atp Binding Residues Of A Protein From Its Primary Sequence. Bmc Bioinformatics, 10: 434.

Coelho, P. S., Bryan, A. C., Kumar, A., Shadel, G. S., And M. Snyder. 2002. A Novel Mitochondrial Protein, Tar1p, Is Encoded On The Antisense Strand Of The Nuclear 25s Rdna. Genes & Development, 16(21), 2755–2760.

Docampo, R., Huang, G. 2015. Calcium Signaling In Trypanosomatid Parasites. Cell Calcium, 57, Pp. 194–202

Grabherr, M.G., Haas, B.J., Yassour, M., Levin, J.Z., Thompson, D.A., Amit, I., Adiconis, X., Fan, L., Raychowdhury, R., Zeng, Q., Chen, Z., Mauceli, E., Hacohen, N., Gnirke, A., Rhind, N., Di Palma, F., Birren, B.W., Nusbaum C., Lindblad-Toh, K., Friedman, N., And A. Regev. 2011. Full-Length Transcriptome Assembly From Rna-Seq Data Without A Reference Genome. Nat Biotechnol. 15:644–52.

Gelder, S.R. 1980. A Review Of The Symbiotic Oligochaeta (Annelida). Zoologischer Anzeiger Jena 204: 69–81.

Glennon, E.K.K., Austin, L.S., Arang, N., Kain, H.S., Mast, F.D., Vijayan K., Aitchison Jd., Kappe Shi., And A. Kaushansky. Alterations In Phosphorylation Of Hepatocyte Ribosomal Protein S6 Control Plasmodium Liver Stage Infection. Cell Rep. 2019 Mar 19;26(12):3391–3399.E4. Doi: 10.1016/J.Celrep.2019.02.085. Pmid: 30893610; Pmcid: Pmc6447308.

Goodchild., C.G. 1951. A New Endoparasitic Oligochaete (Naididae) From A North American Tree-Toad, Hyla Squirella Latreille, 1802. Journal Of Parasitology 37: 205–211.

Haas, B.J., Papanicolaou, A., Yassour, M., Grabherr, M., Blood, P.D., Bowden, J., Couger, M.B., Eccles, D., Li B., Lieber, M., Macmane, S M.D., Ott M., Orvis, J., Pochet, N., Strozz, I F., Weeks, N., Westerman, R., William, T., Dewey, C.N., Henschel, R., Leduc, R.D., Friedman, N., And A. Regev. 2013. De Novo Transcript Sequence Reconstruction From Rna-Seq Using The Trinity Platform For Reference Generation And Analysis. Nat Protoc. 8:1494–512.

Harman, W. J. And A.R. Lawler. 1975. Dero (Allodero) Hylae, An Oligochaete Symbiont In Hylid Frogs In Mississippi. Transactions Of The American Microscopical Society, 94(1), 38–42.

Harman, W.J. 1973. Dero (Allodero) Hylae (Oligochaeta: Naididae) In Louisiana Anurans. Proceedings Of The Louisiana Academy Of Sciences 36: 71–76.

Ingram, G.M. 1997. Analysis Of Cell Cycle Associated Proteins From Theileria Annulata. Retrieved April 16, 2022, From Https://Theses.Gla.Ac.Uk/74753/

Iwami, M. 2000. Bombyxin:An Insect Brain Peptide That Belongs To The Insulin Family. Zoolog Science.17:1035–44.

Kawabe, Y., Waterson, H., And A. Mizoguchi. 2019. Bombyxin (Bombyx Insulin-Like Peptide) Increases The Respiration Rate Through Facilitation Of Carbohydrate Catabolism In *Bombyx Mori*. Front. Endocrinol. 10:150. Doi: 10.3389/Fendo.2019.00150

Langeland, A., Hawdon, J.M., And D.M. O’halloran. 2021. Nemchr-Db: A Database Of Parasitic Nematode Chemosensory G-Protein Coupled Receptors. 51: 333 – 337.

Leszczynski, J., And M.K. Shukla. 2014. Nucleic Acids: Ground-State And Excited-State Properties, Structures, And Interactions And Environmental Aspects As Revealed By Computational Studies, Reference Module In Chemistry, Molecular Sciences And Chemical Engineering, Elsevier

Manni, M., Berkeley, M.R., Seppey, M., And E.M. Zdobnov. 2021. Busco: Assessing Genomic Data Quality And Beyond. Current Protocols, 1, E323.

Michaelsen, W. 1926. Schmarotzende Oligochaten Nebst Erorterungen Uber Verwandt-Schaftliche Beziehungen Der Archioligochaten. Mitt. Zool. St. Inst. Zool. Mus. Ham-Burg, 42: 91–103.

Nagamune, K., Moreno, S.N., Chini, E.N., And L.D. Sibley. 2008. Calcium Regulation And Signaling In Apicomplexan Parasites. In: Burleigh, B.A., Soldati-Favre, D. (Eds) Molecular Mechanisms Of Parasite Invasion. Subcellular Biochemistry, Vol 47. Springer, New York, Ny.

Nwosu, Z.C., Ebert, M.P., Dooley, S. And C. Meyer. 2016. Caveolin-1 In The Regulation Of Cell Metabolism: A Cancer Perspective. Molecular Cancer 15: 71.

Roberts, L. S., And J. Janovy. 2008. Foundations Of Parasitology, 8th Ed. Mcgraw-Hill, Boston, Massachusetts, 728 P.

Robinson, M.D., Mccarthy, D.J., And G.K. Smyth. 2010. Edger: A Bioconductor Package For Differential Expression Analysis Of Digital Gene Expression Data. Bioinformatics 26: 139–140.

Sauvage, V., Aubert, D., Escotte-Binet, S. And I. Villena 2009. The Role Of Atp-Binding Cassette (Abc) Proteins In Protozoan Parasites Mol. Biochem. Parasitol., 167: 81–94.

Sperber, C. 1948. A Taxonomical Study Of The Naididae. Zool. Bidrag Fran Uppsala, 28: 1–296

Uniprot Consortium, T. (2018). Uniprot: The Universal Protein Knowledgebase. Nucleic Acids Research, 46(5), 2699–2699.

